# How cells tame noise while maintaining ultrasensitive transcriptional responses

**DOI:** 10.1101/2025.06.12.659288

**Authors:** Eui Min Jeong, Chang Yoon Chung, Jae Kyoung Kim

**Author notes:** Corresponding author: ∼4. Department of Mathematical Sciences, KAIST, 291, Daehak-ro Yuseong-gu, Daejeon 34141 Republic of Korea (Jae Kyoung Kim). These authors contributed equally to this work (Eui Min Jeong, Chang Yoon Chung).

## Abstract

Ultrasensitive transcriptional switches are essential for converting gradual molecular inputs into decisive gene expression responses, enabling critical behaviors such as bistability and oscillations. While cooperative binding, relying on direct repressor-DNA binding, has been classically regarded as a key ultrasensitivity mechanism, recent theoretical works have demonstrated that combinations of indirect repression mechanisms—sequestration, blocking, and displacement—can also achieve ultrasensitive switches with greater robustness to transcriptional noise. However, these previous works have neglected key biological constraints such as DNA binding kinetics and the limited availability of transcriptional activators, raising the question of whether ultrasensitivity and noise robustness can be sustained under biologically realistic conditions. Here, we systematically assess the impact of these factors on ultrasensitivity and noise robustness under physiologically plausible conditions. We show that while various repression combinations can reduce noise, only the full combination of all three indirect mechanisms consistently maintains low noise and high ultrasensitivity. As a result, biological oscillators employing this triple repression architecture retain precise rhythmic switching even under high noise, and even when activators are shared across thousands of target genes. Our findings offer a mechanistic explanation for the frequent co-occurrence of these repression mechanisms in natural gene regulatory systems.

**AUTHOR SUMMARY:** Cells must make accurate decisions in noisy environments using limited molecular resources. One essential tool for this is the ultrasensitive transcriptional switch, which enables sharp transitions in gene expression. While cooperative binding has long been viewed as the primary mechanism behind ultrasensitivity, it is highly sensitive to molecular noise. Our study explores how cells overcome this limitation by combining three indirect repression mechanisms: sequestration, blocking, and displacement. We show that this triple-repression architecture not only generates ultrasensitivity but also ensures noise robustness under physiologically realistic conditions—even when a single pool of transcription factors regulates thousands of genes. These findings reveal a biologically feasible strategy for noise-resilient transcription and offer a mechanistic explanation for why these repression strategies frequently co-occur in natural systems.

## INTRODUCTION

Ultrasensitivity, characterized by a sharp output change in response to a small variation in input, underlies many essential regulatory functions in biological systems, including signal amplification, bistability, and oscillatory dynamics [1–4]. One of the most well-known mechanisms that produce ultrasensitive responses is cooperative binding, in which transcriptional repressors bind cooperatively to multiple sites on DNA. This mechanism enables the formation of sharp transcriptional switches, where small changes in repressor concentration lead to abrupt transcriptional repression [5–9]. While a cooperative binding mechanism has been traditionally considered a primary strategy for achieving ultrasensitivity, recent theoretical studies have highlighted alternative approaches [5, 10–17]. In particular, combinations of indirect repression mechanisms—such as sequestration, blocking, and displacement—have been shown through ordinary differential equation (ODE) models to produce comparably strong ultrasensitive responses without requiring direct DNA binding [18]. Building on this, stochastic modeling studies further revealed that unlike cooperative binding, which is susceptible to transcriptional noise, systems employing multiple indirect repression mechanisms exhibit enhanced noise robustness [19]. Collectively, these findings suggest that combinations of indirect repression mechanisms can generate ultrasensitive transcriptional switches that are more robust to noise than a cooperative binding mechanism.

However, these previous studies typically assumed fixed or idealized conditions, without explicitly accounting for key biological parameters such as DNA binding/unbinding rates and the abundance of transcriptional activators [18, 19]. In living cells, the kinetics of molecular interactions and the limited availability of regulatory proteins impose significant constraints on transcriptional dynamics [20–22]. It remains unclear whether ultrasensitivity and noise robustness can still be achieved under such physiologically realistic conditions. Moreover, it is unknown which combinations of repression mechanisms, if any, are capable of maintaining performance under these constraints. Addressing this gap is crucial for understanding how biological systems maintain both precision and sensitivity in gene regulation despite molecular noise and finite resources.

To address this, we explicitly incorporated DNA binding rates and activator copy numbers into the models and systematically evaluated how these biologically constrained parameters impact ultrasensitivity and noise robustness. Our results show that, although various indirect repression combinations can reduce noise when DNA binding is fast and activators are abundant, only the full combination of all three mechanisms consistently achieves low noise levels and high ultrasensitivity within physiologically plausible constraints. Accordingly, biological oscillators employing this triple repression architecture can generate precise rhythms, reliably toggling the transcriptional switch on and off even in the presence of stochastic fluctuations. We further demonstrate that this robustness is preserved even when a single pool of activators simultaneously regulates thousands of target genes rather than a single gene, highlighting the scalability and efficiency of this architecture under conditions mimicking a natural gene regulatory network. These findings offer a mechanistic rationale for the frequent co-occurrence of sequestration, blocking, and displacement in natural transcriptional circuits and present a design principle for constructing resource-efficient and noise-resilient gene regulatory systems.

## RESULTS

### Ultrasensitivity generated with cooperative binding is noisy

The cooperative binding mechanism, in which the transcriptional repressors bind cooperatively to multiple DNA sites to inhibit transcription, is one of the most common transcriptional mechanisms for generating ultrasensitivity [5–9]. To achieve an ultrasensitive transcriptional response, we used a model describing the cooperative binding with four independent binding sites (Fig. 1a). In the model, free DNA (*E*_0000_) contains four binding sites, where repressors (*R*) bind at a rate of *k*_*f*_ and unbind with rates of *k*_*r*_, *ck*_*r*_, *c*^2^*k*_*r*_, and *c*^3^*k*_*r*_, for cases in which one (*E*_0001_, *E*_0010_, *E*_0100_, and *E*_1000_), two (*E*_1100_, *E*_1010_, *E*_1001_, *E*_0110_, *E*_0101_, and *E*_0011_), three (*E*_1110_, *E*_1101_, *E*_1011_, and *E*_0111_), and all four (*E*_1111_) sites are occupied by *R*, respectively. When all sites are occupied, transcription is inhibited, but it remains active when at least one site is unoccupied, producing mRNA at a rate of *⍺* , which degrades at a rate of *β* (Table 1). Thus, the transcriptional activity (i.e., the probability at which the transcription is active) decreases as the number of repressors (*R*_*T*_) increases. Notably, the transcriptional activity can be changed sensitively with respect to changes in *R*_*T*_ in the presence of cooperativity (i.e., *c* < 1), achieving ultrasensitivity comparable to the Hill exponent of four (Fig. 1b). This ultrasensitivity is maintained as long as the dissociation constants *K*_*r*_ = *k*_*r*_/*k*_*f*_ remains fixed, even when *k*_*f*_varies. However, the noise level changes when *k*_*f*_ varies. For instance, as *k*_*f*_ = 6 × 10^2^ is increased to *k*_*f*_ = 6 × 10^5^, while keeping *K*_*r*_ = 10^-2^, the level of ultrasenstivity does not change (Fig. 1b), but the noise level quantified with the Fano factor (i.e., variance/mean) of mRNA copy numbers considerably decreases (Fig. 1c). To investigate how much faster *k*_*f*_ and *k*_*r*_ attenuate transcriptional noise, we simulated the stationary distribution of mRNA copy numbers (Fig. 1d). When *k*_*f*_ and *k*_*r*_became slow relative to the rates of mRNA production and degradation (i.e., *⍺* and *β*), slow transitions between the active and repressed DNA states resulted in bimodal mRNA stationary distributions (Fig. 1d, top). Such a bimodality deviates the mRNA distribution from a Poisson distribution, leading to high Fano factor. In contrast, when binding and unbinding rates were faster, the mRNA distribution became unimodal, closely resembling a Poisson distribution which has the Fano factor of one (Fig. 1d, bottom).

**Fig. 1:**
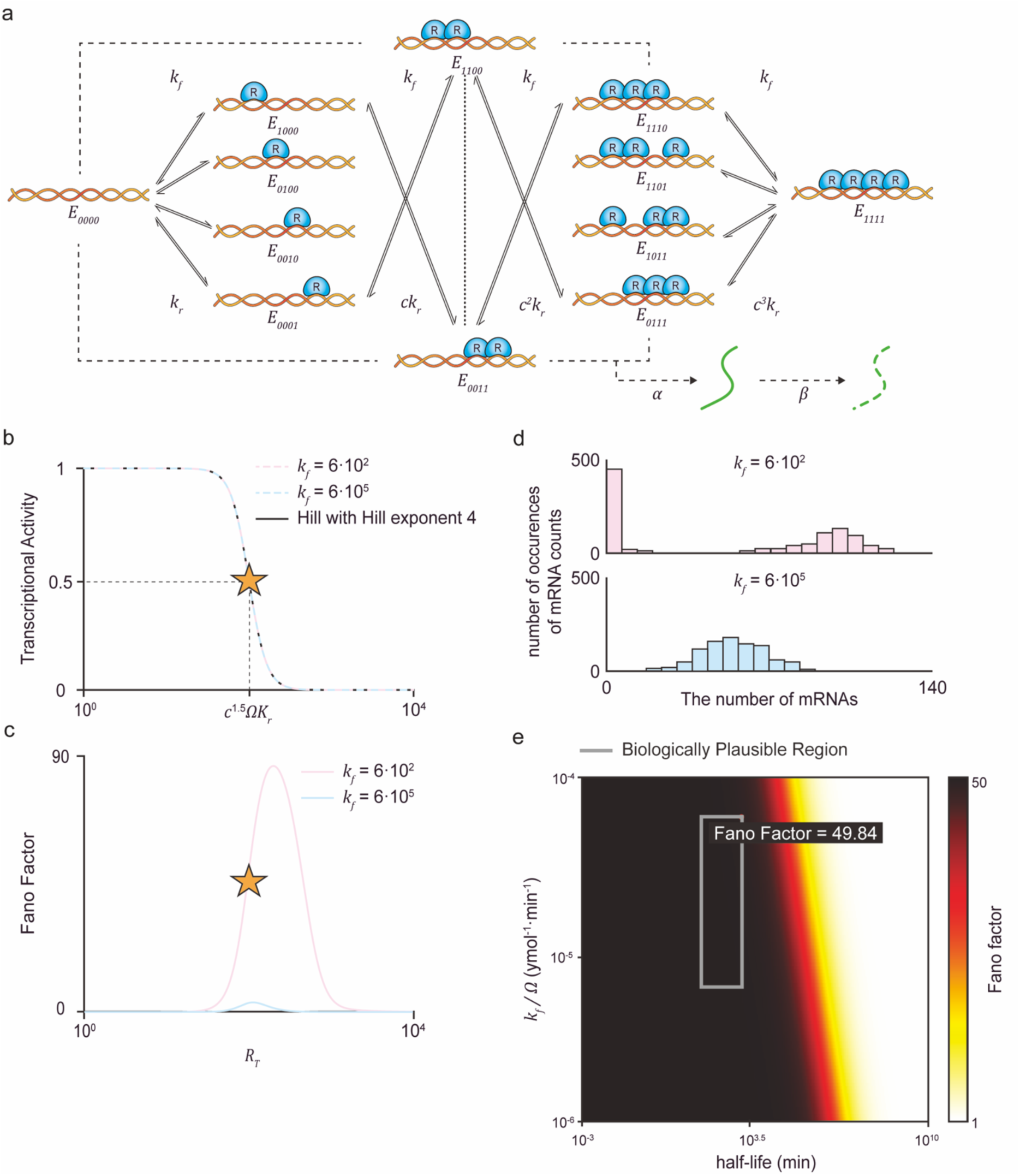
The cooperative binding mechanism is sensitive to noise under biologically realistic conditions. (a) Model diagram of the transcription regulated by the repressor proteins (*R*) binding to four independent sites on the DNA within a cell volume of Ω. Each binding site is occupied by *R* at a rate *k*_*f*_. Conversely, *R* unbinds from DNA at a rate *k*_*r*_when one site is occupied, with the dissociation constant between *R* and DNA defined as *K*_*r*_ = *k*_*r*_/*k*_*f*_. For two, three, or four occupied sites, *R* unbinds at rates of *ck*_*r*_, *c*^2^*k*_*r*_, and *c*^3^*k*_*r*_, respectively. Accordingly, when *c* < 1, cooperative binding is present. When all binding sites are occupied, transcription is inhibited. On the other hand, when any sites remain unoccupied, mRNA is produced at a rate *⍺* and degrades at a rate *β*. (b) When *c* = 10^-4^, transcriptional activity closely resembles the Hill equation with Hill exponent of 4 (black line). Furthermore, transcriptional activity remains consistent with the Hill function regardless of *k*_*f*_ values, provided *K*_*r*_ is kept constant (blue dashed line and red dashed line). (c) Nevertheless, the overall noise level quantified by the Fano factor of mRNAs shows significant differences based on *k*_*f*_. Such differences are particularly evident when transcriptional activity undergoes sensitive response. Notably, the overall noise levels are reduced during the sensitive response when *k*_*f*_is faster (red solid line) compared to when it is slower (blue solid line). (d) To investigate how differences in *k*_*f*_ values affect noise levels, the stationary distributions of mRNAs are simulated at the transcriptional activity of 0.5, where transcriptional activity exhibits the most sensitive response (b and c, star). A slower *k*_*f*_ results in DNA dynamics that are slower relative to mRNA dynamics (i.e., *k*_*f*_Ω^-1^, *k*_*r*_ ≪ *⍺*, *β*), causing slow transitions between inhibited and activated DNA complexes. These slow transitions between inhibited and activated DNA complexes lead to a bimodal distribution of mRNAs that significantly deviates from the Poisson distribution, leading to a high Fano factor (top). On the other hand, a faster *k*_*f*_ produces unimodal distributions that resemble a Poisson distribution, causing a reduced Fano factor (bottom). (e) Since the relative rates of DNA dynamics (i.e., *k*_*f*_Ω^-1^ and *k*_*r*_) and mRNA dynamics (i.e., *⍺* and *β*) affect noise levels, the Fano factors of mRNA distributions are examined under varying rates. Specifically, the Fano factor at the transcriptional activity of 0.5 is calculated with respect to *k*_*f*_ and the half-life of mRNA (ln 2 /*β*), while maintaining the dissociation constant *K*_*r*_ = *k*_*r*_/*k*_*f*_ and effective transcription rate *⍺*/*β*. When *k*_*f*_or the half-life is higher and thus *k*_*f*_becomes faster relative to mRNA dynamics, the Fano factor decreases. However, with biologically realistic *k*_*f*_and half-life (gray box), noise levels remain substantially high (Fano factor > 45).

**Table 1.**
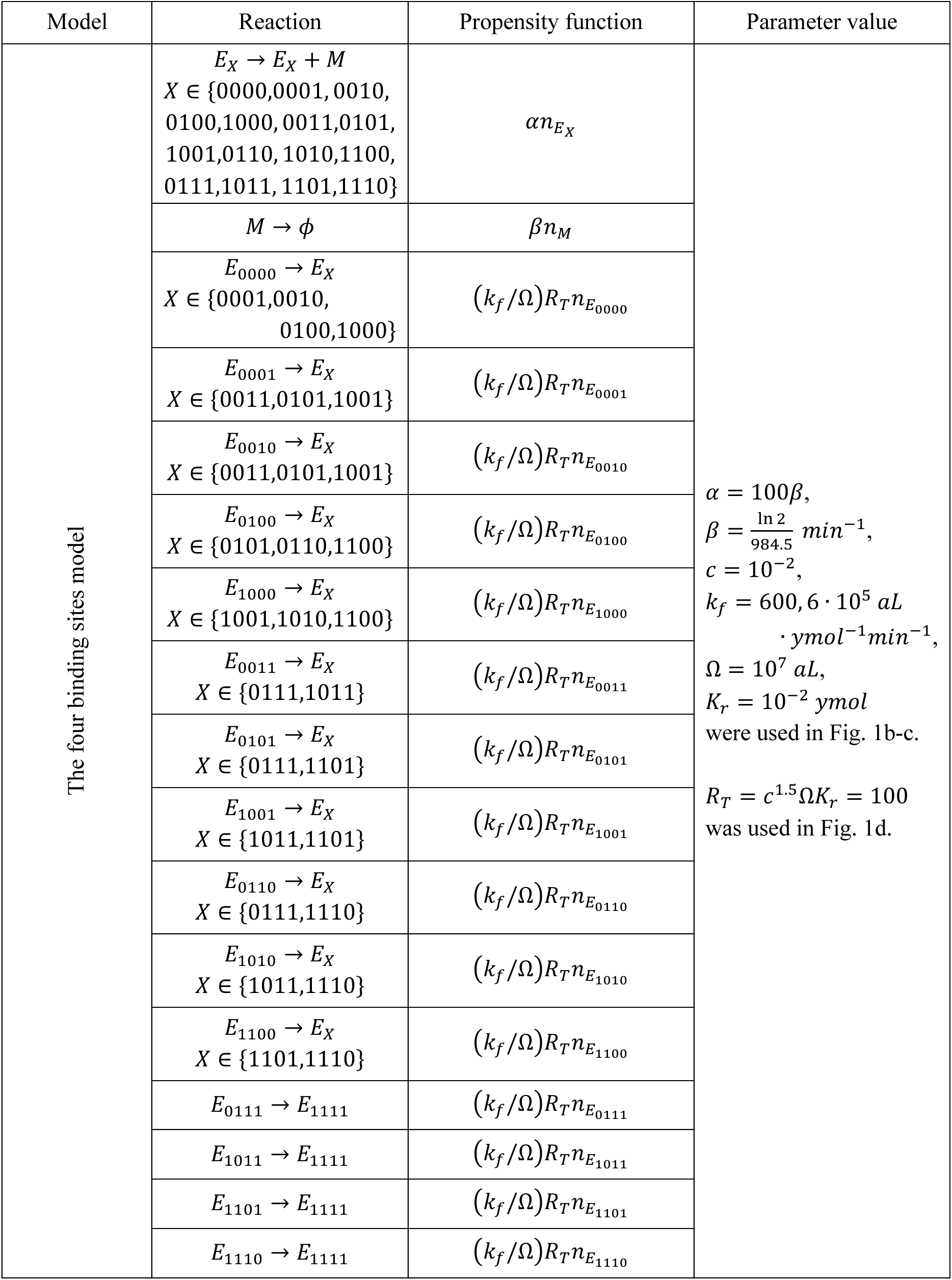
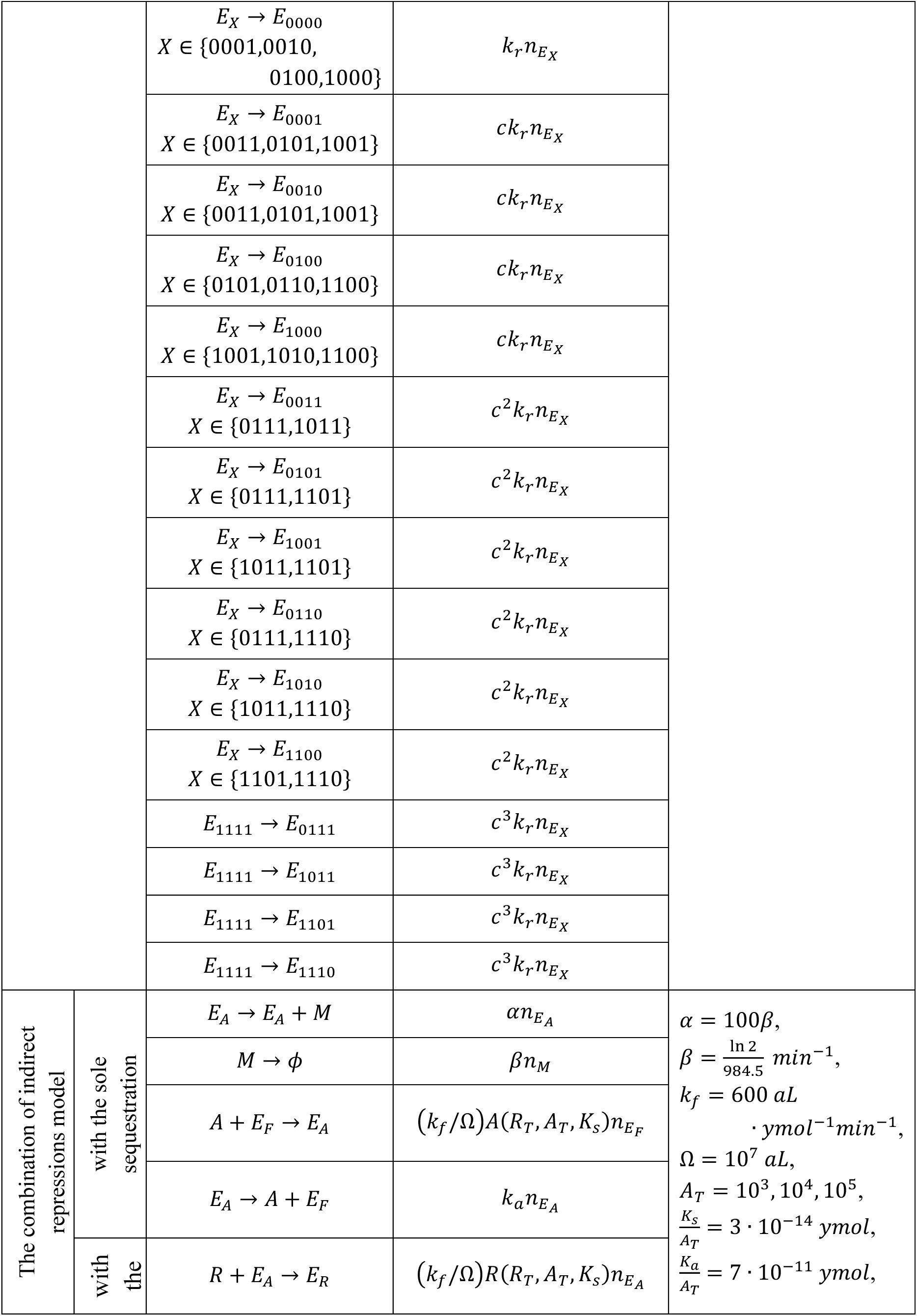
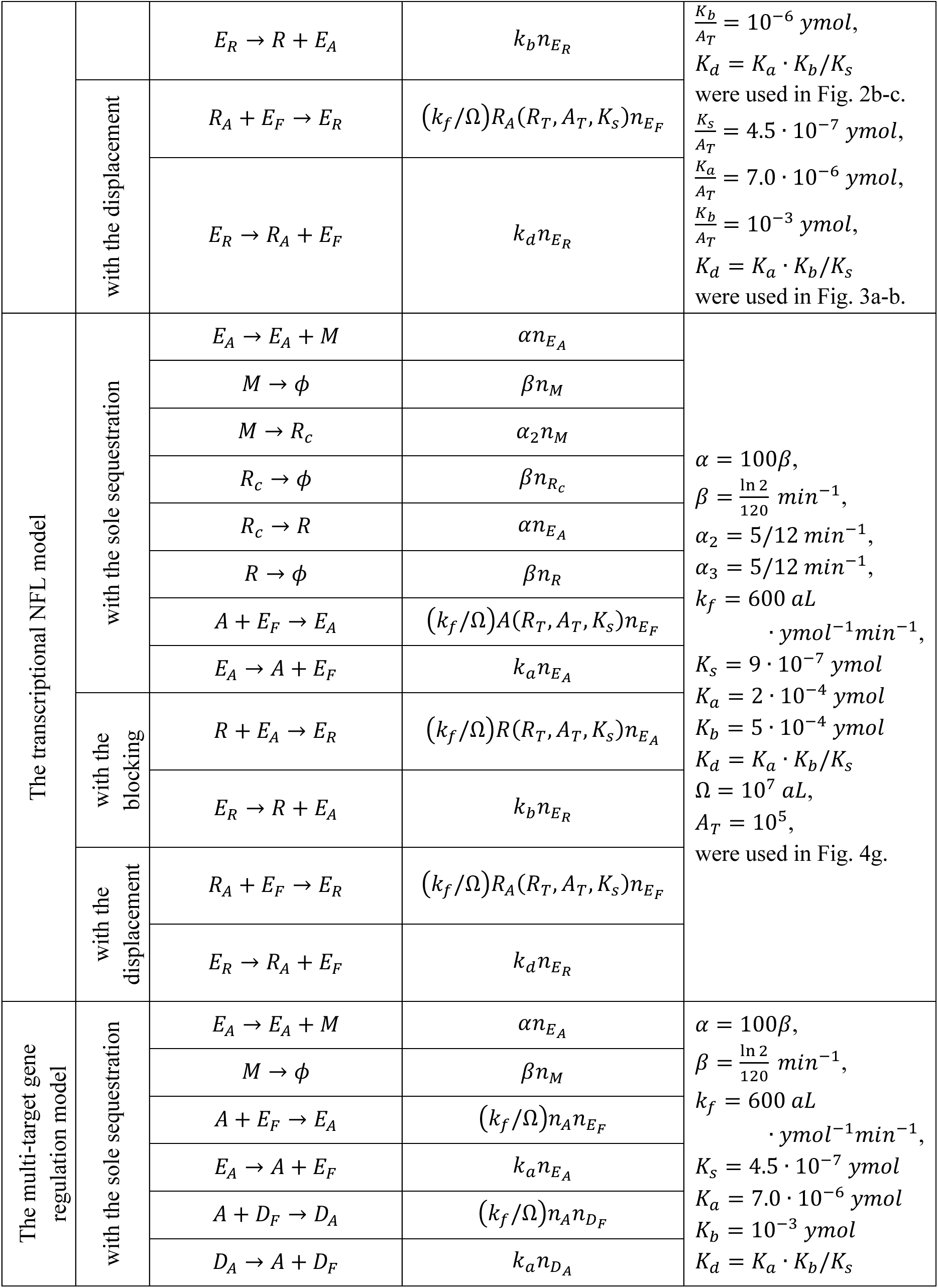
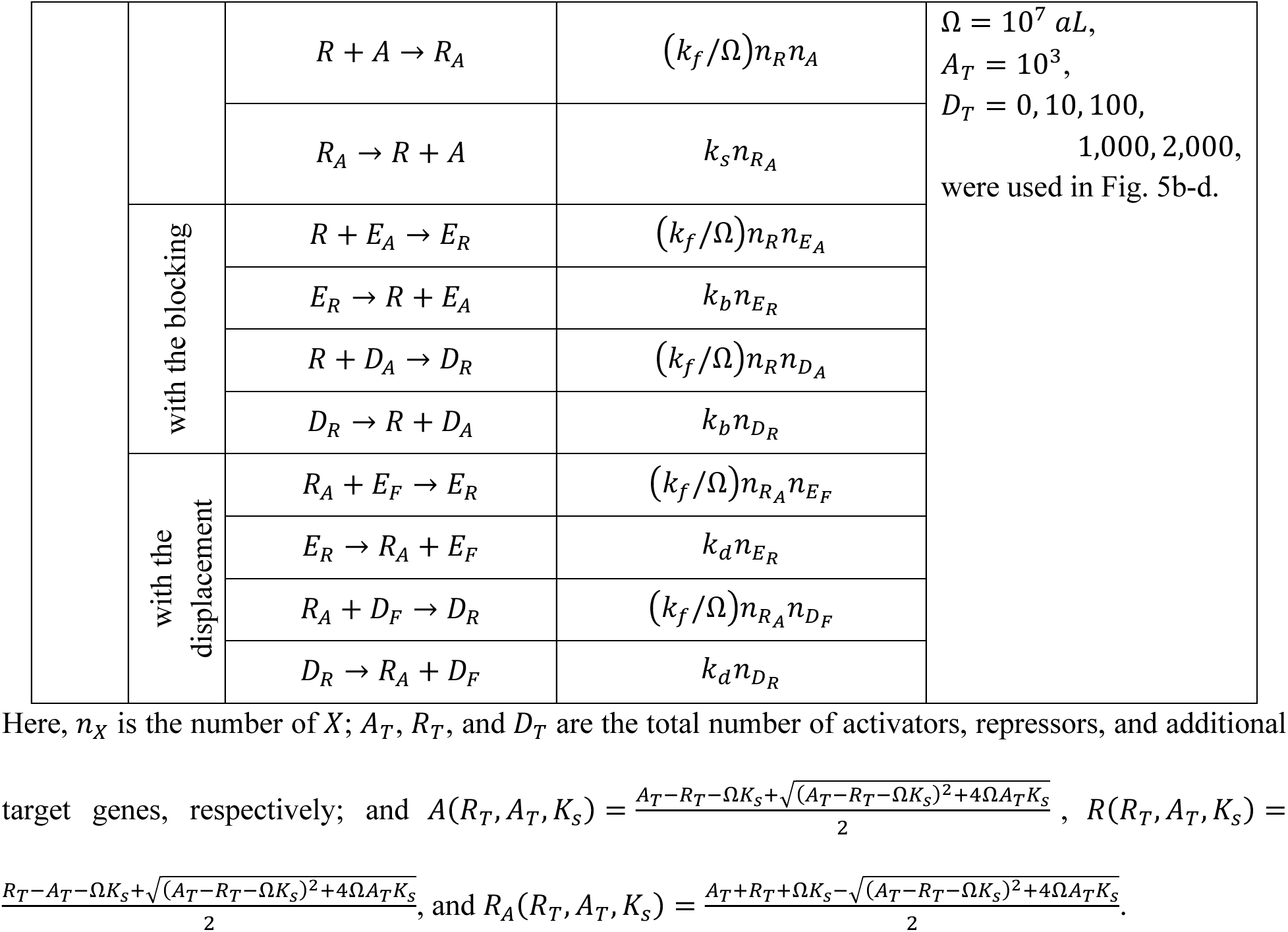
Propensity functions of reactions and parameter values for all models.

| Model | Reaction | Propensity function | Parameter value |
| --- | --- | --- | --- |
| The four binding sites model | $E_X \rightarrow E_X + M$<br>$X \in \{0000,0001,0010,0100,1000,0011,0101,1001,0110,1010,1100,0111,1011,1101,1110\}$ | $\alpha n_{E_X}$ | $\alpha = 100\beta,$<br>$\beta = \frac{\ln 2}{984.5} \text{ min}^{-1},$<br>$c = 10^{-2},$<br>$k_f = 600, 6 \cdot 10^5 \text{ aL}$<br>$\cdot \text{ ymol}^{-1} \text{ min}^{-1},$<br>$\Omega = 10^7 \text{ aL},$<br>$K_r = 10^{-2} \text{ ymol}$<br>were used in Fig. 1b-c.<br>$R_T = c^{1.5} \Omega K_r = 100$<br>was used in Fig. 1d. |
| | $M \rightarrow \phi$ | $\beta n_M$ | |
| | $E_{0000} \rightarrow E_X$<br>$X \in \{0001,0010,0100,1000\}$ | $(k_f/\Omega) R_T n_{E_{0000}}$ | |
| | $E_{0001} \rightarrow E_X$<br>$X \in \{0011,0101,1001\}$ | $(k_f/\Omega) R_T n_{E_{0001}}$ | |
| | $E_{0010} \rightarrow E_X$<br>$X \in \{0011,0101,1001\}$ | $(k_f/\Omega) R_T n_{E_{0010}}$ | |
| | $E_{0100} \rightarrow E_X$<br>$X \in \{0101,0110,1100\}$ | $(k_f/\Omega) R_T n_{E_{0100}}$ | |
| | $E_{1000} \rightarrow E_X$<br>$X \in \{1001,1010,1100\}$ | $(k_f/\Omega) R_T n_{E_{1000}}$ | |
| | $E_{0011} \rightarrow E_X$<br>$X \in \{0111,1011\}$ | $(k_f/\Omega) R_T n_{E_{0011}}$ | |
| | $E_{0101} \rightarrow E_X$<br>$X \in \{0111,1101\}$ | $(k_f/\Omega) R_T n_{E_{0101}}$ | |
| | $E_{1001} \rightarrow E_X$<br>$X \in \{1011,1101\}$ | $(k_f/\Omega) R_T n_{E_{1001}}$ | |
| | $E_{0110} \rightarrow E_X$<br>$X \in \{0111,1110\}$ | $(k_f/\Omega) R_T n_{E_{0110}}$ | |
| | $E_{1010} \rightarrow E_X$<br>$X \in \{1011,1110\}$ | $(k_f/\Omega) R_T n_{E_{1010}}$ | |
| | $E_{1100} \rightarrow E_X$<br>$X \in \{1101,1110\}$ | $(k_f/\Omega) R_T n_{E_{1100}}$ | |
| | $E_{0111} \rightarrow E_{1111}$ | $(k_f/\Omega) R_T n_{E_{0111}}$ | |
| | $E_{1011} \rightarrow E_{1111}$ | $(k_f/\Omega) R_T n_{E_{1011}}$ | |
| | $E_{1101} \rightarrow E_{1111}$ | $(k_f/\Omega) R_T n_{E_{1101}}$ | |
| | $E_{1110} \rightarrow E_{1111}$ | $(k_f/\Omega) R_T n_{E_{1110}}$ | |

|  |  |  |  |  |
| --- | --- | --- | --- | --- |
| | | $E_X \rightarrow E_{0000}$<br>$X \in \{0001, 0010, 0100, 1000\}$ | $k_r n_{E_X}$ | |
| | | $E_X \rightarrow E_{0001}$<br>$X \in \{0011, 0101, 1001\}$ | $ck_r n_{E_X}$ | |
| | | $E_X \rightarrow E_{0010}$<br>$X \in \{0011, 0101, 1001\}$ | $ck_r n_{E_X}$ | |
| | | $E_X \rightarrow E_{0100}$<br>$X \in \{0101, 0110, 1100\}$ | $ck_r n_{E_X}$ | |
| | | $E_X \rightarrow E_{1000}$<br>$X \in \{1001, 1010, 1100\}$ | $ck_r n_{E_X}$ | |
| | | $E_X \rightarrow E_{0011}$<br>$X \in \{0111, 1011\}$ | $c^2 k_r n_{E_X}$ | |
| | | $E_X \rightarrow E_{0101}$<br>$X \in \{0111, 1101\}$ | $c^2 k_r n_{E_X}$ | |
| | | $E_X \rightarrow E_{1001}$<br>$X \in \{1011, 1101\}$ | $c^2 k_r n_{E_X}$ | |
| | | $E_X \rightarrow E_{0110}$<br>$X \in \{0111, 1110\}$ | $c^2 k_r n_{E_X}$ | |
| | | $E_X \rightarrow E_{1010}$<br>$X \in \{1011, 1110\}$ | $c^2 k_r n_{E_X}$ | |
| | | $E_X \rightarrow E_{1100}$<br>$X \in \{1101, 1110\}$ | $c^2 k_r n_{E_X}$ | |
| | | $E_{1111} \rightarrow E_{0111}$ | $c^3 k_r n_{E_X}$ | |
| | | $E_{1111} \rightarrow E_{1011}$ | $c^3 k_r n_{E_X}$ | |
| | | $E_{1111} \rightarrow E_{1101}$ | $c^3 k_r n_{E_X}$ | |
| | | $E_{1111} \rightarrow E_{1110}$ | $c^3 k_r n_{E_X}$ | |
| The combination of indirect repressions model | with the sole sequestration | $E_A \rightarrow E_A + M$ | $\alpha n_{E_A}$ | $\alpha = 100\beta,$<br>$\beta = \frac{\ln 2}{984.5} \text{ min}^{-1},$<br>$k_f = 600 \text{ aL}$<br>$\cdot \text{ ymol}^{-1} \text{ min}^{-1},$<br>$\Omega = 10^7 \text{ aL},$<br>$A_T = 10^3, 10^4, 10^5,$<br>$\frac{K_S}{A_T} = 3 \cdot 10^{-14} \text{ ymol},$<br>$\frac{K_a}{A_T} = 7 \cdot 10^{-11} \text{ ymol},$ |
| | | $M \rightarrow \phi$ | $\beta n_M$ | |
| | | $A + E_F \rightarrow E_A$ | $(k_f/\Omega)A(R_T, A_T, K_S)n_{E_F}$ | |
| | | $E_A \rightarrow A + E_F$ | $k_a n_{E_A}$ | |
| | with the | $R + E_A \rightarrow E_R$ | $(k_f/\Omega)R(R_T, A_T, K_S)n_{E_A}$ | |
| | | $E_R \rightarrow R + E_A$ | $k_b n_{E_R}$ | $\frac{K_b}{A_T} = 10^{-6} \text{ ymol},$<br>$K_d = K_a \cdot K_b / K_s$ |
| | with the displacement | $R_A + E_F \rightarrow E_R$ | $(k_f / \Omega) R_A (R_T, A_T, K_s) n_{E_F}$ | were used in Fig. 2b-c.<br>$\frac{K_s}{A_T} = 4.5 \cdot 10^{-7} \text{ ymol},$ |
| | | $E_R \rightarrow R_A + E_F$ | $k_d n_{E_R}$ | $\frac{K_a}{A_T} = 7.0 \cdot 10^{-6} \text{ ymol},$<br>$\frac{K_b}{A_T} = 10^{-3} \text{ ymol},$<br>$K_d = K_a \cdot K_b / K_s$<br>were used in Fig. 3a-b. |
| The transcriptional NFL model | with the sole sequestration | $E_A \rightarrow E_A + M$ | $\alpha n_{E_A}$ | $\alpha = 100\beta,$<br>$\beta = \frac{\ln 2}{120} \text{ min}^{-1},$<br>$\alpha_2 = 5/12 \text{ min}^{-1},$<br>$\alpha_3 = 5/12 \text{ min}^{-1},$<br>$k_f = 600 \text{ aL}$<br>$\cdot \text{ ymol}^{-1} \text{ min}^{-1},$<br>$K_s = 9 \cdot 10^{-7} \text{ ymol}$<br>$K_a = 2 \cdot 10^{-4} \text{ ymol}$<br>$K_b = 5 \cdot 10^{-4} \text{ ymol}$<br>$K_d = K_a \cdot K_b / K_s$<br>$\Omega = 10^7 \text{ aL},$<br>$A_T = 10^5,$<br>were used in Fig. 4g. |
| | | $M \rightarrow \phi$ | $\beta n_M$ | |
| | | $M \rightarrow R_c$ | $\alpha_2 n_M$ | |
| | | $R_c \rightarrow \phi$ | $\beta n_{R_c}$ | |
| | | $R_c \rightarrow R$ | $\alpha n_{E_A}$ | |
| | | $R \rightarrow \phi$ | $\beta n_R$ | |
| | | $A + E_F \rightarrow E_A$ | $(k_f / \Omega) A (R_T, A_T, K_s) n_{E_F}$ | |
| | | $E_A \rightarrow A + E_F$ | $k_a n_{E_A}$ | |
| | with the blocking | $R + E_A \rightarrow E_R$ | $(k_f / \Omega) R (R_T, A_T, K_s) n_{E_A}$ | |
| | | $E_R \rightarrow R + E_A$ | $k_b n_{E_R}$ | |
| | with the displacement | $R_A + E_F \rightarrow E_R$ | $(k_f / \Omega) R_A (R_T, A_T, K_s) n_{E_F}$ | |
| | | $E_R \rightarrow R_A + E_F$ | $k_d n_{E_R}$ | |
| The multi-target gene regulation model | with the sole sequestration | $E_A \rightarrow E_A + M$ | $\alpha n_{E_A}$ | $\alpha = 100\beta,$<br>$\beta = \frac{\ln 2}{120} \text{ min}^{-1},$<br>$k_f = 600 \text{ aL}$<br>$\cdot \text{ ymol}^{-1} \text{ min}^{-1},$<br>$K_s = 4.5 \cdot 10^{-7} \text{ ymol}$<br>$K_a = 7.0 \cdot 10^{-6} \text{ ymol}$<br>$K_b = 10^{-3} \text{ ymol}$<br>$K_d = K_a \cdot K_b / K_s$ |
| | | $M \rightarrow \phi$ | $\beta n_M$ | |
| | | $A + E_F \rightarrow E_A$ | $(k_f / \Omega) n_A n_{E_F}$ | |
| | | $E_A \rightarrow A + E_F$ | $k_a n_{E_A}$ | |
| | | $A + D_F \rightarrow D_A$ | $(k_f / \Omega) n_A n_{D_F}$ | |
| | | $D_A \rightarrow A + D_F$ | $k_a n_{D_A}$ | |
| | | $R + A \rightarrow R_A$ | $(k_f/\Omega)n_R n_A$ | $\Omega = 10^7 \text{ aL},$<br>$A_T = 10^3,$<br>$D_T = 0, 10, 100,$<br>$1,000, 2,000,$<br>were used in Fig. 5b-d. |
| | | $R_A \rightarrow R + A$ | $k_s n_{R_A}$ | |
| | with the blocking | $R + E_A \rightarrow E_R$ | $(k_f/\Omega)n_R n_{E_A}$ | |
| | | $E_R \rightarrow R + E_A$ | $k_b n_{E_R}$ | |
| | | $R + D_A \rightarrow D_R$ | $(k_f/\Omega)n_R n_{D_A}$ | |
| | | $D_R \rightarrow R + D_A$ | $k_b n_{D_R}$ | |
| | with the displacement | $R_A + E_F \rightarrow E_R$ | $(k_f/\Omega)n_{R_A} n_{E_F}$ | |
| | | $E_R \rightarrow R_A + E_F$ | $k_d n_{E_R}$ | |
| | | $R_A + D_F \rightarrow D_R$ | $(k_f/\Omega)n_{R_A} n_{D_F}$ | |
| | | $D_R \rightarrow R_A + D_F$ | $k_d n_{D_R}$ | |

Here, $n_X$ is the number of $X$ ; $A_T$ , $R_T$ , and $D_T$ are the total number of activators, repressors, and additional
target genes, respectively; and $A(R_T, A_T, K_S) = \frac{A_T - R_T - \Omega K_S + \sqrt{(A_T - R_T - \Omega K_S)^2 + 4\Omega A_T K_S}}{2}$ , $R(R_T, A_T, K_S) =$
$\frac{R_T - A_T - \Omega K_S + \sqrt{(A_T - R_T - \Omega K_S)^2 + 4\Omega A_T K_S}}{2}$ , and $R_A(R_T, A_T, K_S) = \frac{A_T + R_T + \Omega K_S - \sqrt{(A_T - R_T - \Omega K_S)^2 + 4\Omega A_T K_S}}{2}$ .

Next, we investigate whether the noise level in transcription decreases as the binding and unbinding rates become faster across a broad parameter range. For this, while keeping *K*_*r*_ = 10^-2^ and *⍺*/*β* = 100 , we varied *k*_*f*_ and *β*. We found that, when the binding rate is sufficiently fast relative to the mRNA half-life (ln 2 /*β*), the Fano factor approached its minimal value of 1 (Fig. 1e). However, achieving this requires biologically unrealistic fast binding and unbinding rates. Specifically, with a biologically relevant binding rate *k*_*f*_Ω^-1^ = 6 × 10^-6^∼6 × 10^-5^ *ymol*^-1^*min*^-1^[21], where Ω = 10^7^*aL* is the volume of the cell [20], and the mRNA half-life is 0.5 ∼ 16.4 hours (i.e., 30 ∼ 984 min) [23, 24], the Fano factor is greater than 40 (Fig. 1e, gray box). Taken together, while increasing binding rates can minimize noise, the cooperative binding mechanism remains highly noisy under physiological conditions.

### Ultrasensitivity generated with indirect repression mechanisms is robust to noise

The transcriptional repression by repressors can occur not only through direct binding to DNA such as cooperative binding mechanism, but also by indirectly inhibiting activators that promote transcription [12, 25–28]. In addition, combining multiple indirect repression mechanisms can generate an ultrasensitive transcriptional response [18], while maintaining lower noise levels [19]. Accordingly, we investigated whether the models describing different combinations of indirect repression mechanisms can achieve ultrasensitivity with low transcriptional noise under biologically plausible conditions.

In this model, activators (*A*) bind to free DNA (*E*_*F*_) at a rate *k*_*f*_ to form activator-bound DNA (*E*_*A*_), and unbind at a rate *k*_*a*_. Once *E*_*A*_is formed, the transcription is activated, leading to mRNAs production at a rate *⍺* and degradation at a rate *β* (Fig. 2a, bottom). To inhibit the transcription, indirect repression mechanisms such as sequestration, blocking, and displacement can be used. Specifically, free repressors (*R*) can bind to *A* at a rate *k*_*f*_, forming an activator-repressor complex (*R*_*A*_) to prevent *A* from binding to DNA (sequestration; Fig. 2a, top left), while unbinding from *R*_*A*_ at a rate *k*_*s*_. *R* can also bind directly to *E*_*A*_ at a rate *k*_*f*_, forming the activator-repressor-bound DNA (*E*_*R*_), thereby blocking transcription (blocking; Fig. 2a, top middle), while *R* can unbind from *E*_*R*_ at a rate *k*_*b*_. Additionally, *R* removes *A* from *E*_*A*_ with the form of *R*_*A*_ at a rate *k*_*d*_, whereas *R*_*A*_ can bind to DNA at a rate *k*_*f*_ reversely (displacement; Fig. 2a, top right, and Table 1). Based on these reactions, we constructed three distinct models one with sole sequestration by including only *A* , *R* , and *R*_*A*_dynamics; a second with sequestration and blocking by additionally incorporating *R* binding/unbinding to *E*_*A*_; and a third model with all three repressions by further including binding/unbinding between *R*_*A*_ and *E*_*F*_.

**Fig. 2:**
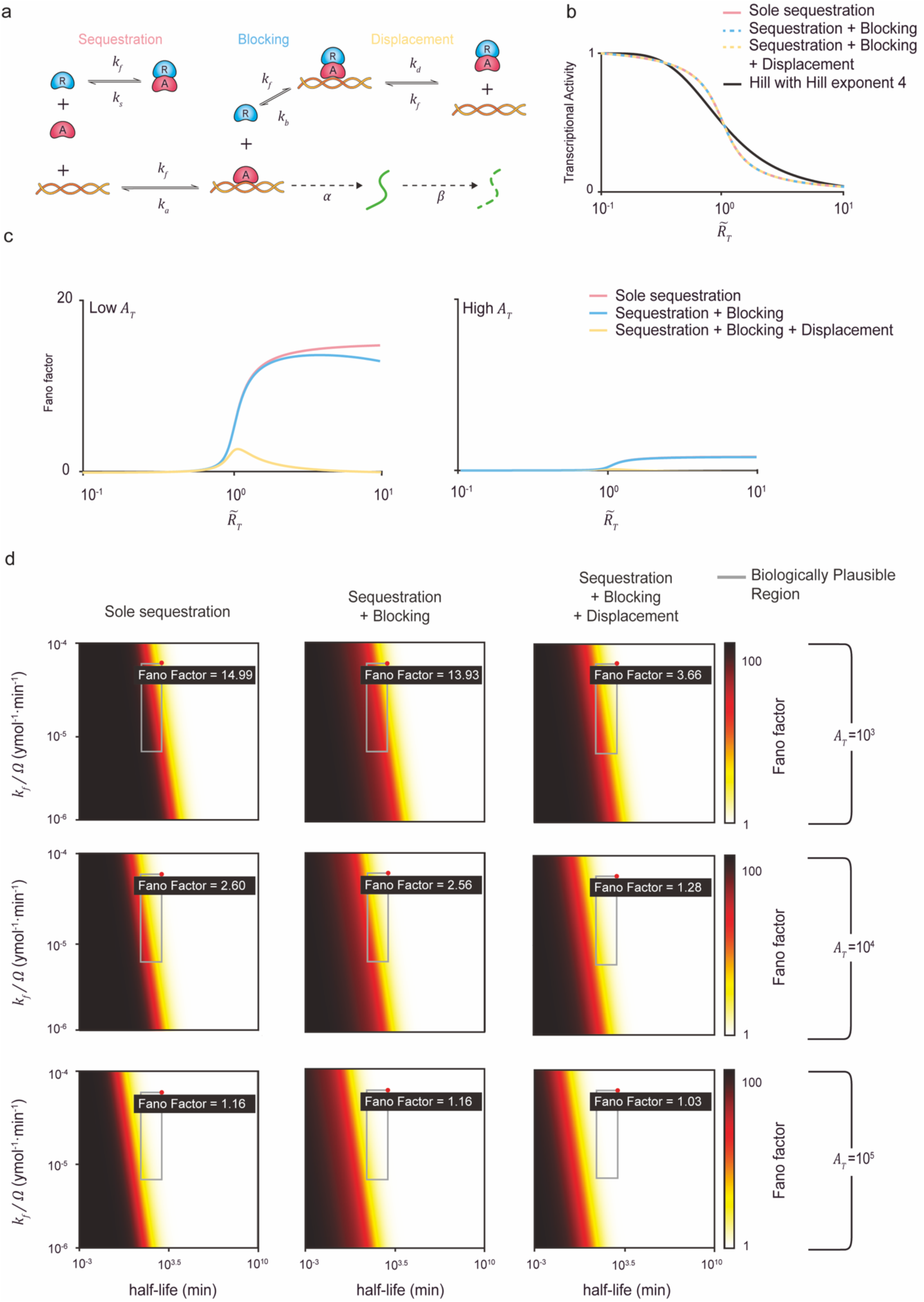
The combinations of repression mechanisms can achieve minimal noise levels with high sensitivity under biologically realistic conditions. (a) Model diagram of transcription regulated by multiple repression mechanisms. An activator (*A*) binds to DNA, forming an activated DNA complex (*E*_*A*_) at a rate *k*_*f*_, and unbinds from DNA at a rate *k*_*a*_. Once *E*_*A*_ is formed, mRNAs are produced at a rate *⍺* and degrade at a rate *β*. To prevent the formation of *E*_*A*_, a repressor (*R*) sequesters free *A* at a rate *k*_*f*_, forming a repressor-activator complex (*R*_*A*_; sequestration). *R* also unbinds from *R*_*A*_at a rate *k*_*s*_. When *A* is already bound to DNA, a free *R* binds directly to the DNA-bound *A* at a rate *k*_*f*_ forming a repressed DNA complex (*E*_*R*_; blocking), and unbinds from *E*_*R*_at a rate *k*_*b*_. Additionally, free *R* can displace DNA-bound *A* from *E*_*R*_ by forming *R*_*A*_ (displacement) at a rate *k*_*d*_, while *R*_*A*_ binds to DNA at a rate *k*_*fE*_. Accordingly, the dissociation constants are defined as *K*_*a*_ = *k*_*a*_/*k*_*f*_, *K*_*s*_ = *k*_*s*_/*k*_*f*_, *K*_*b*_ = *k*_*b*_/*k*_*f*_, and *K*_*d*_ = *k*_*d*_/*k*_*f*_. (b) To investigate whether the transcriptional activity of multiple repression mechanisms can exhibit a sensitive response similar to the Hill function, transcriptional activities of the sole sequestration (red solid line), the combination of sequestration and blocking (blue dashed line), and the combination of sequestration, blocking, and displacement (orange dashed line) are derived. For all three models, when the number of repressors (*R*_*T*_) is less than the number of activators (*A*_*T*_), unsequestered activators can bind to DNA, promoting transcription (activation phase). However, when *R*_*T*_ exceeds *A*_*T*_ , most activators are sequestered by repressors, thereby suppressing transcription (repression phase). In the switching from activation to repression phase, a sharp transition occurs when the molar ratio between *A*_*T*_ and *R*_*T*_ (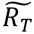 = *R*_*T*_/*A*_*T*_) is near one. The sensitivity of this transition is determined by the dissociation constants *K*_*a*_, *K*_*s*_, *K*_*b*_, and *K*_*d*_; hence, they are appropriately adjusted to match all models that exhibit transcriptional activity comparable to the Hill function with Hill exponent of 4 (black solid line). (c) Despite this similarity in transcriptional activity, the three models show significant differences in overall noise levels. Specifically, overall noise levels decrease with the addition of repression mechanisms: from the sole sequestration (red solid line) to the combination of sequestration and blocking (blue solid line), and further the combination of sequestration, blocking, and displacement (orange solid line). Additionally, overall noise levels across all 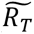 values are lower when *A*_*T*_ is higher (right) compared to when it is lower (left). (d) Given the impact of *A*_*T*_ on overall noise levels across the three models, the Fano factor of mRNAs is examined under varying *A*_*T*_ values. Specifically, the maximum Fano factor near *R*^T^_*T*_ of 1 is calculated with respect to *k*_*f*_Ω^-1^ and the half-life of mRNA, while maintaining the dissociation constant *K*_*a*_, *K*_*b*_ and *K*_*d*_, and effective transcription rate *⍺*/*β*. Heatmaps of the maximum Fano factor are shown for the sole sequestration (first column), the combination of sequestration and blocking (second column), and the combination of sequestration, blocking and displacement (third column). Each model is simulated with *A*_*T*_ values of 10^3^, 10^4^, and 10^5^, all of which are biologically realistic. When *A*_*T*_ = 10^3^(first row), all three models show lower noise levels compared to the cooperative binding model over the same range of *k*_*f*_Ω^-1^and the half-life of mRNA. Furthermore, overall noise levels progressively decrease as the repression mechanisms are added. In particular, within the biologically realistic *k*_*f*_Ω^-1^and the mRNA half-life (gray box), the combination of all three mechanisms reduces the noise level down to the Fano factor of approximately 3, much lower compared to the cooperative binding (i.e., Fano factor = 49). The noise levels are further decreased as the level of *A*_*T*_increases. When *A*_*T*_ = 10^5^ (third row), the minimum noise levels (i.e., a Fano factor of 1) are achievable within the biologically realistic *k*_*f*_ and the mRNA half-life (gray box) for all three mechanisms.

First, by adjusting the dissociation constants *K*_*s*_ = *k*_*s*_/*k*_*f*_ for sequestration, the model with sequestration alone can achieve ultrasensitivity comparable to that of cooperative binding with four binding sites (Fig. 2b, red line). Subsequently, adjusting *K*_*b*_ = *k*_*b*_/*k*_*f*_ for blocking and *K*_*d*_ = *k*_*d*_/*k*_*f*_for displacement enables other two models to reach similar levels of ultrasenstivity (Fig. 2b, blue and yellow lines). While the level of their ultrasensitivity is similar, interestingly, their noise levels differ (Fig. 2c and Table 2). Specifically, as more repression mechanisms are added, the overall noise levels decrease, consistent with a previous study [19].

**Table 2.**
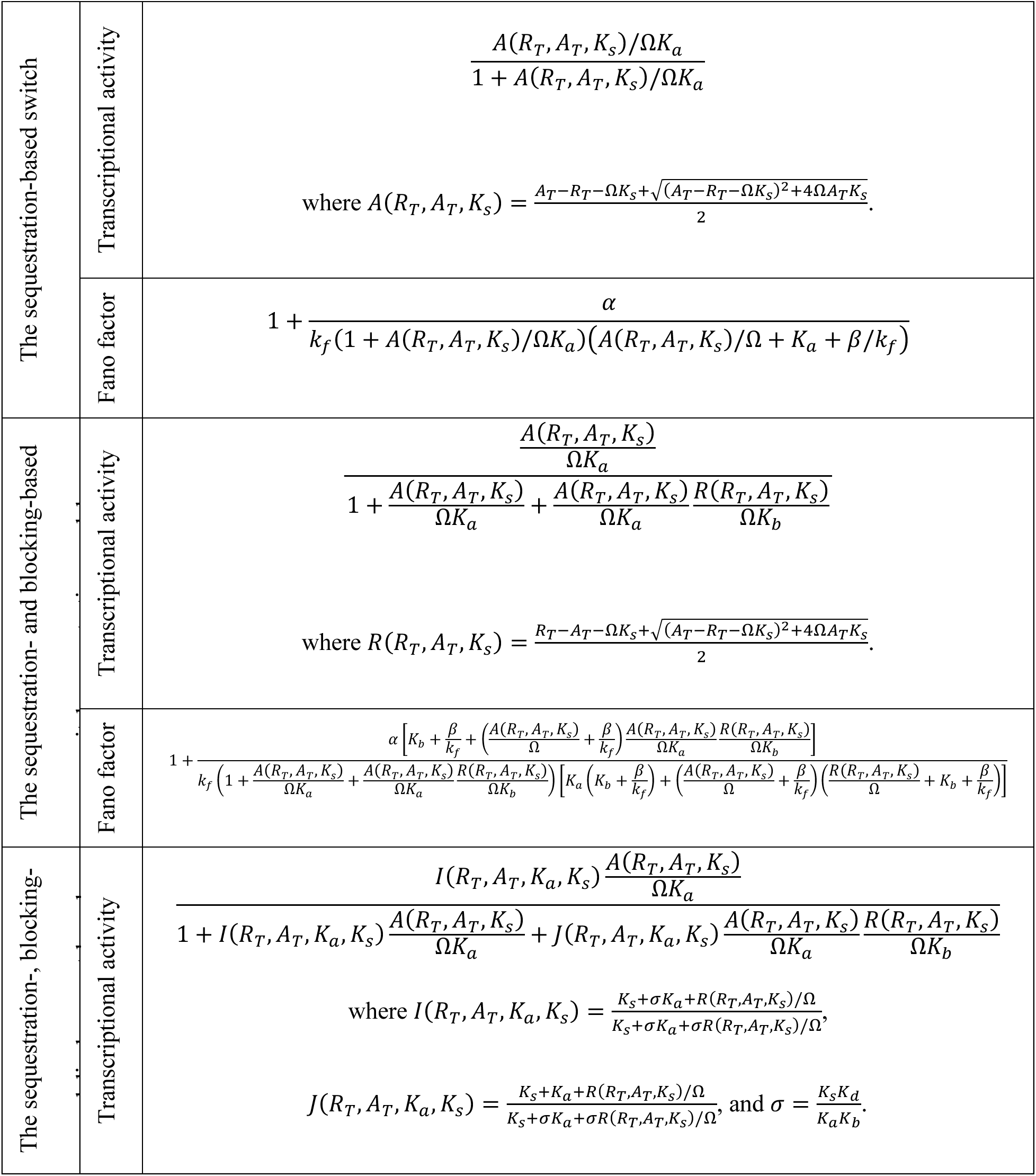
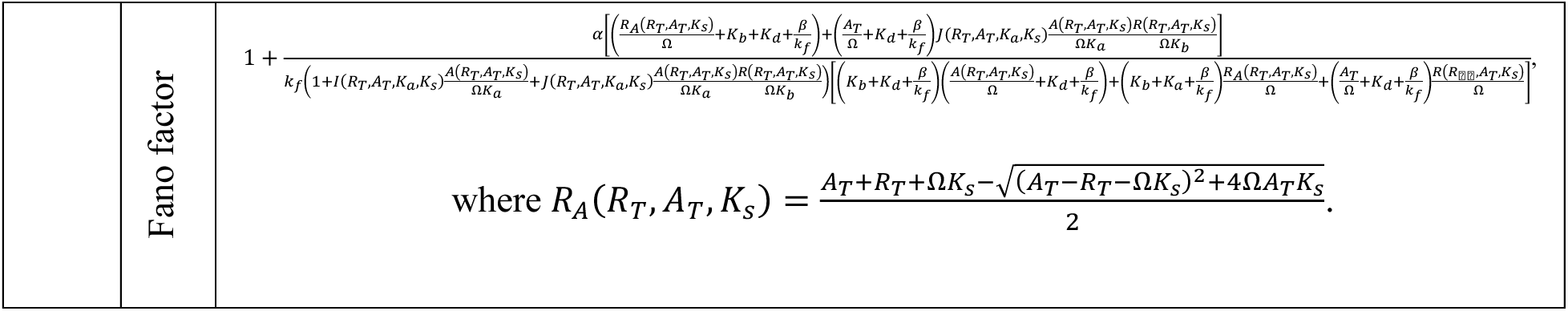
The transcriptional activity and the Fano factor for all models describing combinations of indirect repression mechanisms.

Next, we investigate whether the noise levels can be further decreased as binding and unbinding rates become faster as shown in cooperative binding. Indeed, as binding rates become faster relative to mRNA half-life, the Fano factor decreases in all three models. However, within the biologically relevant range of binding rate and mRNA half-life, the Fano factor still remains high, with a minimum of about 4 (Fig. 2d, top row). In this case, we used the number of activators *A*_*T*_ = 10^3^. Since the biologically relevant copy number of activators ranges from *A*_*T*_ = 10^3^∼10^5^ yoctomole [20], we further increase *A*_*T*_. Indeed, higher *A*_*T*_ shifts the lower noise region closer to the biologically relevant region, without compromising *k*_*f*_ or mRNA half-life (Fig. 2d, middle row). Specifically, high *A*_*T*_ (i.e., *A*_*T*_ = 10^5^) in conjunction with increased binding rate and mRNA half-life can reduce noise levels close to 1 for all three models, without losing biological relevance (Fig. 2d, bottom row).

Importantly, by adjusting the dissociation constants, the combinations of indirect repression mechanisms can generate higher levels of ultrasensitivity (the Hill exponent of 250; Fig. 3a). This elevated ultrasensitivity is accompanied by increased transcriptional noise (Fig. 2c and 3b). In such case, under biologically relevant conditions, only the combination of sequestration, blocking, and displacement is capable of reducing the Fano factor close to 4 (Fig. 3c).

**Fig. 3:**
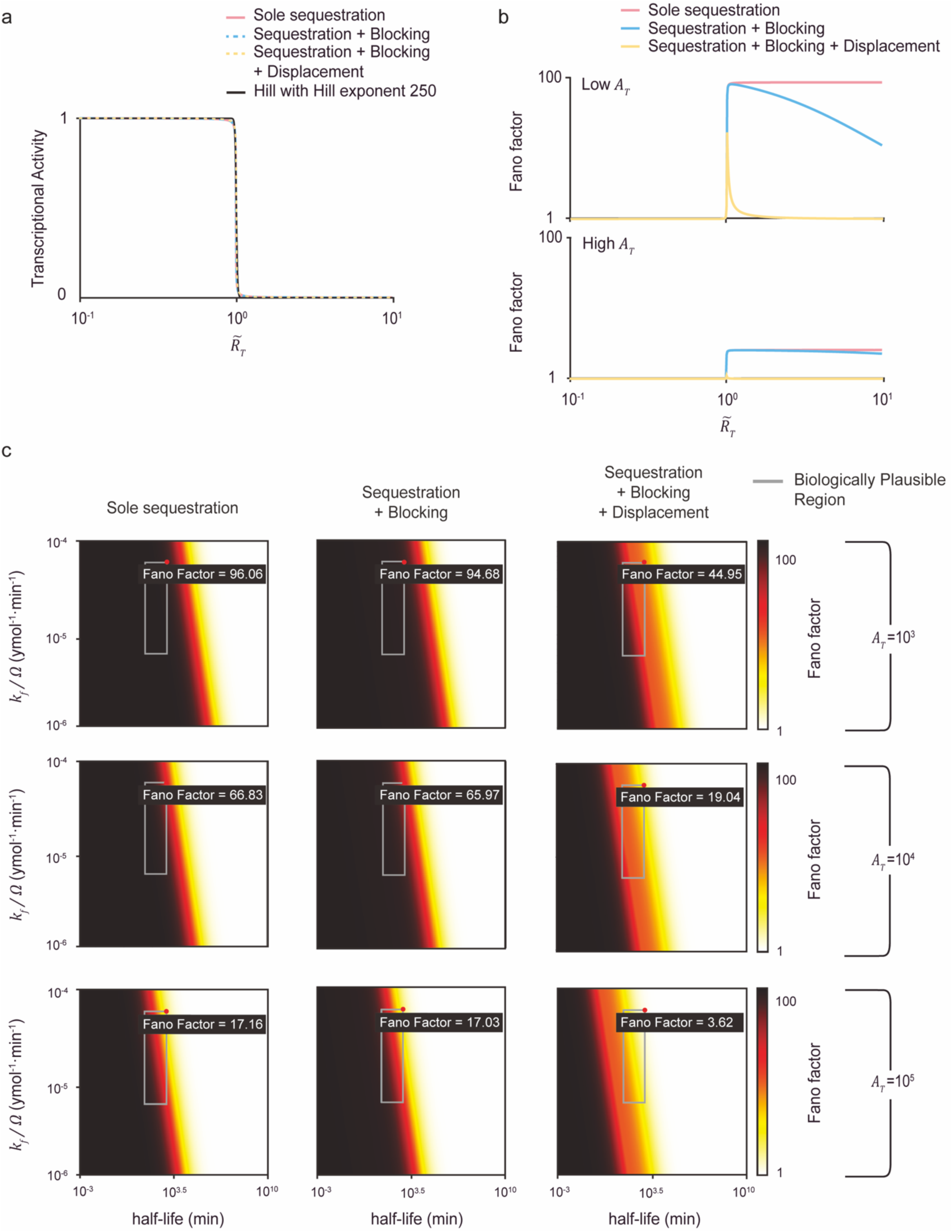
Only the combination of sequestration, blocking and displacement can achieve close to the minimum noise levels with considerably high sensitivity under biologically realistic conditions. (a) The transcriptional activities of the sole sequestration (red solid line), the combination of sequestration and blocking (blue dashed line), and the combination of sequestration, blocking, and displacement (orange dashed line) are matched to resemble the Hill function with Hill exponent of 250 (black solid line) by adjusting the dissociation constants *K*_*a*_, *K*_*s*_, *K*_*b*_, and *K*_*d*_. (b) Despite similarities in transcriptional activities, the three models show significant differences in overall noise levels (top). These noise levels decrease with the addition of repression mechanisms, from the sole sequestration (red solid line) to the combination of sequestration and blocking (blue solid line), and further to the combination of sequestration, blocking, and displacement (orange solid line). Notably, more sensitive responses with Hill exponent of 250 induce greater noise levels compared to those observed with Hill exponent of 4 (Fig. 2c, top). Furthermore, increasing *A*_*T*_leads to an overall reduction in noise levels (bottom). (c) The maximum Fano factor for each model under varying *A*_*T*_values is shown. When *A*_*T*_ = 10^3^ (first row), all models show high noise levels (i.e., Fano factor > 40) over the biological range of *k*_*f*_Ω^-1^and the mRNA half-life. Noise levels decrease as *A*_*T*_ increases (2^nd^ and 3^rd^ rows). In particular, when *A*_*T*_ = 10^5^, the combination of all three mechanisms can lead to reduced noise levels (i.e., a Fano factor close to 3). On the other hand, sole sequestration still results in considerable noise levels (i.e., Fano factor = 17).

### Multiple biological oscillators employ a combination of sequestration, blocking, and displacement mechanisms to produce rhythms that are both robust and precise

The combination of sequestration, blocking, and displacement is a common regulatory mechanism employed across diverse biological oscillators. For example, in the mammalian circadian clock, the PER-CRY complex sequesters CLOCK-BMAL1 as well as displaces it from DNA, while CRY blocks the transcriptional activity of CLOCK-BMAL1 bound to DNA [18, 29–35] (Fig. 4a). Similarly, in the NF-κB oscillator, IκB sequesters NF-κB in the cytoplasm to prevent its activation of transcription. Upon external stimulation, IκB is degraded, allowing NF-κB to translocate to the nucleus, and activate transcription. This activation is suppressed as IκB displaces NF-κB from the DNA [36, 37] (Fig. 4b). In the p53-Mdm2 oscillator, Mdm2 sequesters p53 away from DNA, blocks transcriptional activity of p53, and displaces it from DNA through cooperative interactions with other corepressors [38, 39] (Fig. 4c). Since ultrasensitivity is critical to generate rhythms in biological oscillators [1, 3, 4, 12], we hypothesized that such combination of multiple indirect repressions, yielding a noise-resistant ultrasensitive switch (Fig. 4d, left) [18, 19], can generate strong rhythms through high sensitivity, and maintain precise periodicity through accurate timing of transitions between transcriptional on and off states (Fig. 4d, right).

**Fig. 4:**
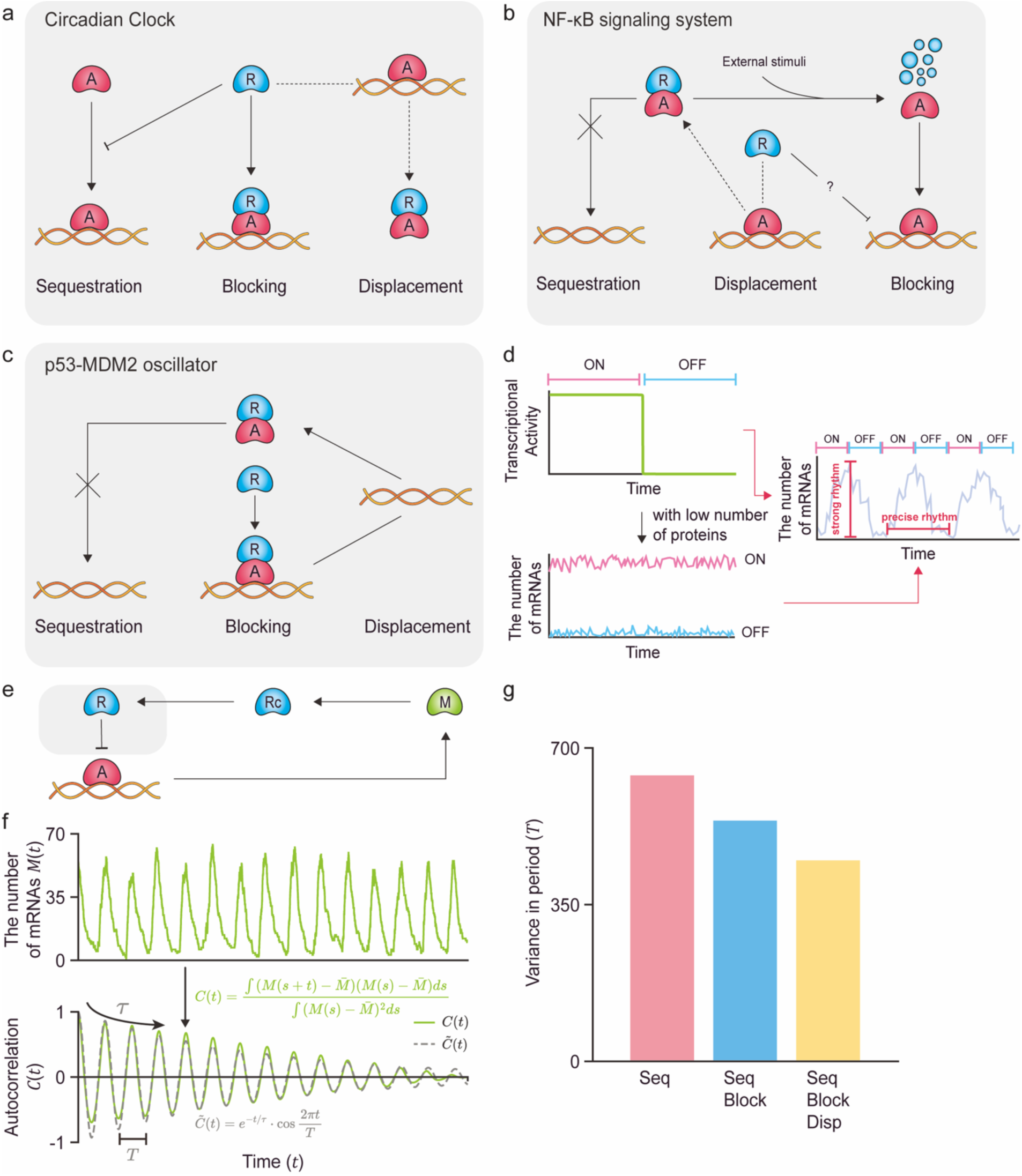
Various biological oscillators utilize the combination of sequestration, blocking, and displacement mechanisms to generate robust and precise rhythms. (a) In the mammalian circadian clock, PER:CRY inhibits CLOCK:BMAL1 by sequestration, preventing its binding to DNA. On the other hand, when CLOCK:BMAL1 is already bound to DNA, CRY blocks transcription or PER:CRY can displace it from DNA. (b) In the NF-κB signaling system, IκB sequesters NF-κB in the cytoplasm, preventing its entry into the nucleus. As IκB degrades in response to external stimuli, the previously sequestered NF-κB is released and enters the nucleus. In the nucleus, NF-κB binds to DNA to promote transcription. IκB can inhibit transcription by displacing NF-κB from DNA, although whether IκB can directly block transcription remains unclear. (c) In the p53-MDM2 oscillator, MDM2 binds to p53, preventing its binding to DNA. On the other hand, when p53 is already bound to DNA, MDM2 inhibits transcription by directly binding to the DNA-bound p53 to block it, as well as displacing it from DNA in cooperation with a corepressor. (d) The combination of sequestration, blocking, and displacement can generate both high sensitivity and robustness against noise, essential for strong and precise rhythms, respectively. Furthermore, even with a low number of proteins (i.e., activators and repressors), mRNAs exhibit low fluctuations in both on and off transcriptional states. Consequently, biological oscillators incorporating all three mechanisms can effectively produce precise rhythms, with accurate increases and decreases during the transcriptional on and off states, respectively. (e) To investigate this, we constructed the transcriptional negative feedback loop (NFL) model. In the NFL model, when the transcription is turned on, mRNA (*M*) is produced, and subsequently translated into the cytoplasmic repressor ( *R*_*c*_). After entering the nucleus, the nucleic repressor ( *R* ) inhibits its own transcription through the multiple repression mechanisms (gray box). (f) Using the NFL model, the oscillatory time-series of mRNA copy numbers (*M*(*t*)) can be simulated (top). To quantify the noise level in *M*(*t*), its autocorrelation function *C*(*t*) (bottom, green solid line) is computed and is fitted with a decaying cosine function 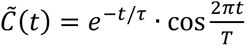 (bottom, gray dashed line). Here, *T* represents the period of oscillation, and *τ* represents how slow *C*(*t*) decays. (g) By simulating *M*(*t*) for three models over 300 times, we evaluated the variance of *T*. As more repression mechanisms are combined, the variance of *T* is reduced, indicating enhanced robustness against noise.

To validate this hypothesis, we constructed a transcriptional negative feedback loop (NFL) model (Fig. 4e and Table 1). In the NFL model, transcriptional activation leads to the synthesis of mRNA (*M*), which is subsequently translated into the repressor in the cytoplasm (*R*_*c*_). Upon nuclear entry, the repressor (*R*) suppresses its own transcription via multiple repression mechanisms (Fig. 4e, gray box). Using this model, we simulated a time-series of mRNA copy numbers (*M*(*t*)) to investigate the robustness of each repression mechanism against noise (Fig. 4f, top). Specifically, we calculated autocorrelation functions of 300 simulated *M*(*t*) (*C*(*t*); Fig. 4f, bottom, green solid line), and fitted them to a decaying cosine function, 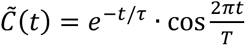, to estimate *T* and *τ* (Fig. 4f, bottom, gray dashed line). By definition of *C̃*(*t*), *T* represents the period of the rhythm, while *τ* reflects how slowly the autocorrelation decays [40, 41]. We observed that, as additional repression mechanisms were incorporated, the variance in the estimated *T* decreased. It implies more consistent periods in each cycle of rhythm under noise (Fig. 4g), indicating enhanced robustness to noise

## DISCUSSION

Previous work demonstrated that combining indirect repression mechanisms—such as sequestration, blocking, and displacement—can generate ultrasensitive transcriptional switches that are more robust to molecular noise than direct mechanisms such as cooperative binding [19]. However, those studies did not account for the timescales of DNA-binding dynamics (i.e., the binding and unbinding rates between DNA and transcriptional factors). In this study, we systematically show that when DNA binding occurs on a faster timescale than mRNA-related processes (i.e., transcription and degradation), ultrasensitivity and noise robustness can be simultaneously achieved in the resulting transcriptional switch (Fig. 1b–e, 2b, and d). We further show that increasing the number of activators can also promote such ultrasensitive behavior (Fig. 2b–d). However, under realistic constraints on binding kinetics and activator abundance, robust ultrasensitive switching is only achieved when all three indirect repression mechanisms are combined (Fig. 3). As a result, biological oscillators incorporating this triple-repression architecture can maintain precise rhythmic transitions between transcriptional states even in the presence of molecular noise (Fig. 4). In summary, our findings suggest that combining indirect repression mechanisms offers a resource-efficient strategy for achieving robust, rhythmic gene expression in noisy cellular environments—potentially explaining why sequestration, blocking, and displacement often co-occur in natural biological oscillators.

In this study, we initially analyzed models in which 1,000 to 100,000 transcriptional activators regulate a single target gene (Fig. 2a). However, in many biological systems employing multiple repression mechanisms—such as the circadian clock, NF-κB oscillator, and p53-Mdm2 oscillator (Fig. 4a–c)— transcriptional activators often regulate hundreds to thousands of genes simultaneously (Fig. 5a). For example, in the mammalian circadian clock, a limited pool of BMAL1 proteins regulates approximately 3,400 genes [42]; p53 controls around 3,700 target genes [43]; and NF-κB targets several hundred genes [44]. To more accurately capture this biological complexity, we extended our model to include varying numbers of target genes (Fig. 5a and Table 1). Remarkably, we found that the transcriptional noise of an individual gene remained largely unaffected, even when the activator pool was shared among thousands of targets (Fig. 5b–d). This result suggests that a finite pool of transcriptional activators can robustly regulate large gene networks without increasing expression noise at the single-gene level, provided that sequestration, blocking, and displacement mechanisms are combined.

**Fig. 5:**
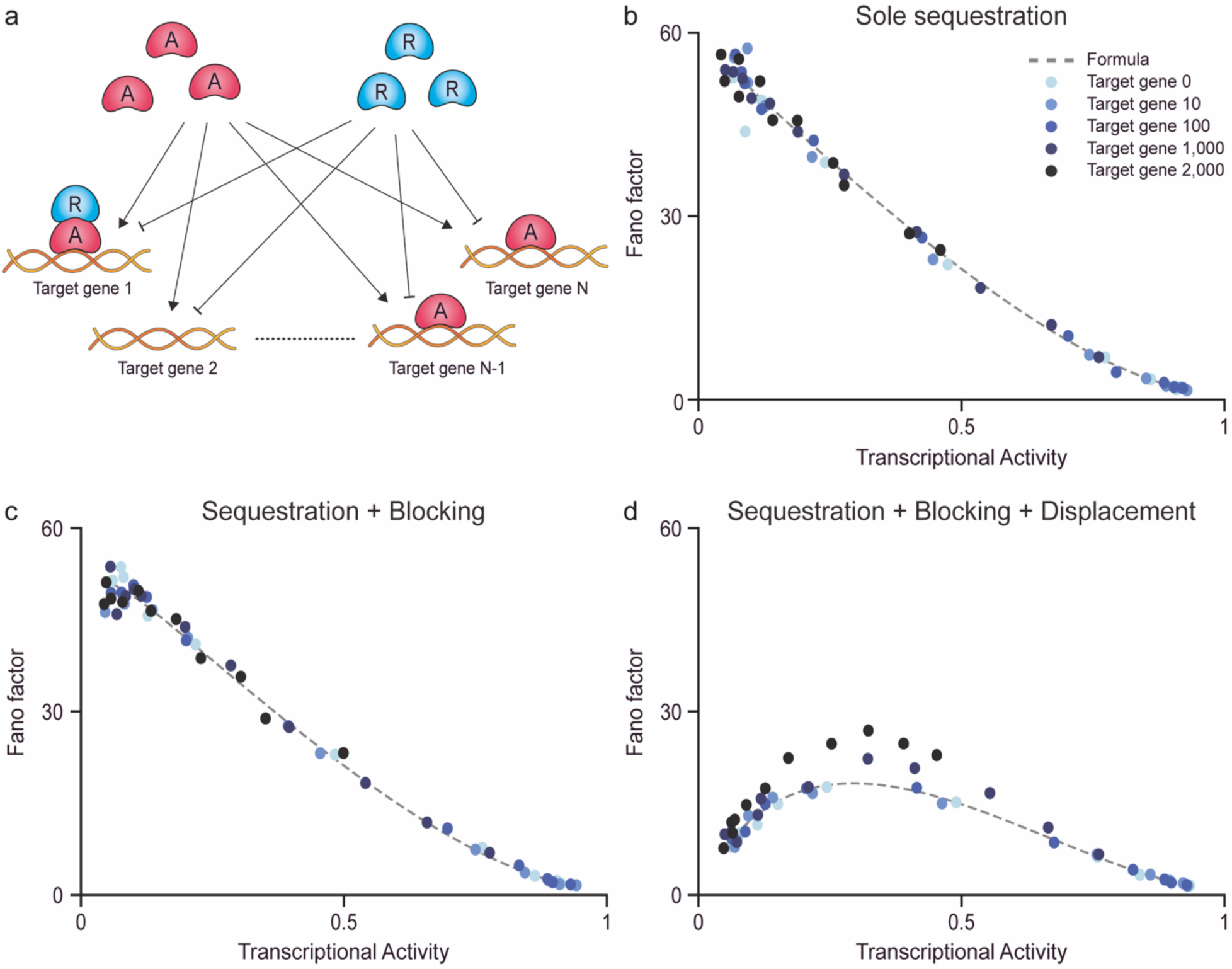
Transcriptional factors regulate thousands of target genes while maintaining noise levels comparable to those of single-gene regulation, given the same level of transcriptional activity. (a) In biological systems, transcriptional activators and repressors often regulate multiple target genes simultaneously. (b–d) We examine whether combining different repression mechanisms can still reduce transcriptional noise. For this, we simulated the stationary distribution of mRNA of one target gene while varying the number of additional target genes to 0, 10, 100, 1,000, and 2,000 (see Methods). From the distribution, we calculated the Fano factor of mRNAs (blue dots). Regardless of the number of target genes, repression via the sole sequestration (b), the combination of sequestration, and blocking (c), the combination of sequestration, blocking, and displacement (d), all show transcriptional noise levels similar to single-gene regulation. Here, dashed lines represent the relationship between the transcriptional activity and the Fano factor without additional target genes.

When the dissociation constants *K*_*a*_, *K*_*s*_, *K*_*b*_, and *K*_*d*_ satisfy the detailed balance condition—i.e., 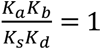 —the binding and unbinding interactions among activators, repressors, and DNA occur at thermodynamic equilibrium, meaning no energy is consumed during transcriptional repression [18]. Under these equilibrium conditions, our results show that simply combining indirect repression mechanisms can enhance the noise robustness of biological oscillators without requiring energy input (Fig. 4). However, recent studies suggest that breaking detailed balance through energy dissipation can further amplify ultrasensitivity. For example, Jeong et al. demonstrated that violating detailed balance in triple-mechanism models yields greater ultrasensitivity than in other combinations [18]. Similarly, Estrada et al. showed that in cooperative binding systems, breaking detailed balance facilitates ultrasensitive responses [45]. Moreover, energy consumption has also been linked to enhanced noise robustness in oscillatory systems: Cao et al. theoretically showed that energy dissipation suppresses phase diffusion due to intrinsic noise in various biochemical oscillators [41], while Fei et al., using the Brusselator model, found that energy expenditure not only reduces phase diffusion but also improves adaptability to environmental changes [46]. Collectively, these findings highlight that beyond combining repression mechanisms, the energy cost of regulatory processes plays a key role in tuning ultrasensitivity and robustness. Investigating how energy dissipation interacts with repression strategies thus presents a compelling direction for future research.

## METHODS

### Derivation of the equations for the transcriptional activity and the Fano factor of the four binding sites model with cooperative binding

To derive the transcriptional activity and Fano factor of a gene regulated by cooperative binding at four DNA sites, we followed the framework introduced by Sanchez et al. [47], formulating a chemical master equation (CME) that captures all 16 possible DNA binding configurations. By computing steady-state moment equations for mRNA distributions, we obtained closed-form expressions for the transcriptional activity and Fano factor as functions of repressor concentration and binding parameters. These derivations quantify how cooperative repression influences gene expression dynamics, and the full mathematical details are provided in the Supplementary Information and at https://github.com/Mathbiomed/Ultrasensitive-gene-switch.

### Derivation of the equations for the transcriptional activity and the Fano factor of the multiple indirect repression mechanism models

Following the approach in the previous section, Jeong et al., derived equations for the transcriptional activity and the Fano factor in models incorporating the multiple indirect repression models [19]. Notably, for analytical convenience, all parameters were normalized by J*k*_*f*_Ω^-1^K*A*_*T*_ . For example, under this normalization, the transcription and degradation rates of mRNA, *⍺* and *β*, were simply switched to *⍺̃* = *⍺*Ω/*k*_*f*_*A*_*T*_ *β̃* and = *β*Ω/*k*_*f*_*A*_*T*_, respectively, and the unbinding rate between activators and DNA, *k*_*a*_, was expressed as the normalized dissociation constant, 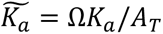. In particular, the binding of activators J*k*_*f*_Ω^-1^K*A*(*R*_*T*_, *A*_*T*_, *K*_*s*_), where the number of free activators is given by

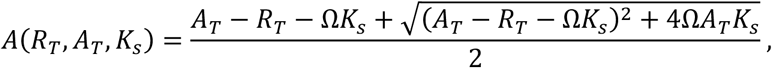

was normalized to the fraction of free activators among the total activators,

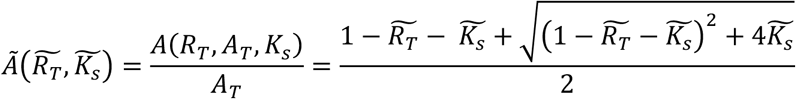

where 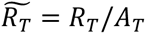 and 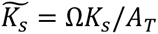 [12, 19, 27, 48–50]. With the normalized variables and parameters, the transcriptional activity and the Fano factor for the sole sequestration model were derived as follows:

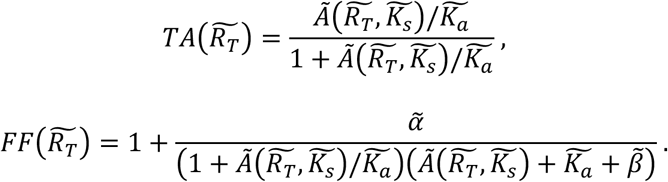

To incorporate *k*_*f*_, *A*_*T*_ , and Ω to the equations explicitly, we substitute, 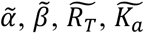 and 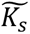 with the original parameters as follows:

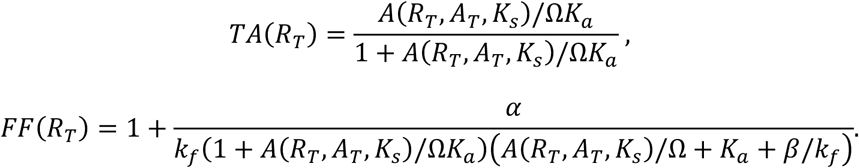

Similarly, we substituted the original parameters into the transcriptional activity and the Fano factor equations derived in Jeong et al for other combinations of indirect repression mechanisms (Table 2).

### Simulations of multi-target genes regulation and noise level quantification

To evaluate how transcriptional noise is affected when transcriptional activators regulate multiple target genes, we extended the multiple indirect repression models to include up to 2,000 co-regulated genes (Fig. 5). To isolate the effect of multi-gene regulation, we fixed the total number of activators to 1,000, while the number of additional target genes was varied from 0 to 2,000. For each gene count, we modulated the transcriptional activity by varying the number of repressors from 0 to 2,000. For each parameter set, we simulated the corresponding CME (Table 1) using the Gillespie algorithm with 1,000 independent runs. The transcriptional activity (i.e., the probability that the gene is active) was quantified as the proportion of runs in which *E*_*A*_ = 1 at the stationarity (Fig. 5b-d, x-axis). Similarly, from resulting stationary mRNA distributions, we calculated the Fano factor to measure the transcriptional noise (Fig. 5b-d, y-axis).

## ACKNOWLEDGEMENTS

The research was supported by the following agencies and institutions: the Institute for Basic Science (grant no. IBS-R029-C3, J.K.K.); Samsung Science and Technology Foundation (grant no. SSTF-BA1902-01, J.K.K.); the National Research Foundation of Korea (NRF) grant funded by the Korean government (MSIT) (no. RS-2022-NR068758).

## SUPPORTING INFORMATION CAPTIONS

S1 Appendix. Supplementary Methods, Derivation of the equations for the transcriptional activity and the Fano factor of the four binding sites model with cooperative binding.

